# Rapid Hippocampal Synaptic Potentiation Induced by Ketamine Metabolite (*2R*,*6R*)-Hydroxynorketamine Persistently Primes Synaptic Plasticity

**DOI:** 10.1101/2024.10.18.619152

**Authors:** Kyle A. Brown, Musa I. Ajibola, Todd D. Gould

## Abstract

The pharmacologically active (*R*,*S*)-ketamine (ketamine) metabolite (*2R*,*6R*)-hydroxynorketamine (HNK) maintains ketamine’s preclinical antidepressant profile without adverse effects. While hypotheses have been proposed to explain how ketamine and its metabolites initiate their antidepressant-relevant effects, it remains unclear how sustained therapeutic actions arise following drug elimination. To distinguish the physiological mechanisms involved in the rapid from sustained actions of HNK, we utilized extracellular electrophysiology combined with pharmacology to develop an *in vitro* hippocampal slice incubation model that exhibited pharmacological fidelity to the 1) rapid synaptic potentiation induced by HNK at the Schaffer collateral-CA1 (SC-CA1) synapse during bath-application to slices collected from mice, and 2) maintenance of metaplastic (priming) activity that lowered the threshold for *N-*methyl-D-aspartate receptor (NMDAR) activation-dependent long-term potentiation (LTP) hours after *in vivo* dosing. We then used this model to reveal novel druggable mechanisms engaged in HNK’s temporally-sensitive antidepressant synaptic actions, finding that the induction of synaptic potentiation by HNK did not require NMDAR activity, but NMDAR activity was necessary to maintain synaptic priming. HNK required protein kinase A (PKA) activity to rapidly potentiate SC-CA1 neurotransmission to facilitate synaptic priming that persistently promoted LTP formation. HNK’s rapid actions were blocked by inhibitors of adenylyl cyclase 1 (AC1), but not an AC5 inhibitor. We conclude that HNK rapidly potentiates SC-CA1 synaptic efficacy, which then stimulates priming mechanisms that persistently favor antidepressant-relevant plasticity. Targeting such priming mechanisms may be an effective antidepressant strategy, and using approaches such as our incubation model may aid in revealing novel pharmacological targets.

## INTRODUCTION

(*R*,*S*)-ketamine (ketamine) alleviates depression symptoms within hours after a subanesthetic infusion in treatment-resistant patients, and relief persists for a day or more after drug elimination [1–3]. Ketamine presents limitations, such as dissociative effects and misuse potential [4], necessitating the development of safer, ketamine-like antidepressants. Synaptic mechanisms leveraged by ketamine to elicit rapid (i.e., <1 hr post-treatment) versus sustained (i.e., hours-days post-treatment) antidepressant effects appear distinct, but converge on amelioration of impaired synaptic strength [5,6]. Yet the precise, time-sensitive, druggable mechanisms ketamine utilizes to evoke antidepressant outcomes remain unclear [7].

Rapid activation of antidepressant synaptic machinery by ketamine is indicated by a “glutamate surge”, which is supported by clinical studies revealing association of antidepressant outcomes with responses indicative of enhanced glutamate release probability in the prefrontal cortex [PFC; 8] and hippocampus [9] during ketamine infusion. Studies in rodents found elevated PFC glutamate levels *ex vivo* [10] and *in vivo* [11,12] minutes after a subanesthetic ketamine dose. Ketamine’s well-recognized pharmacology as an open-channel blocker of the *N*-methyl-D-aspartate receptor (NMDAR) has led to numerous NMDAR inhibition-centered hypotheses to interpret its rapid therapeutic action [13], including disinhibition of glutamatergic neurotransmission following selective inhibition of interneuron NMDARs [14], homeostatic plasticity after transient inhibition of spontaneously active principal cell NMDARs [15], or selectively blocking activity of NMDARs in lateral habenular glutamatergic cells [16–18]. Yet, clinical trials with alternative NMDAR inhibitors have failed to mimic ketamine’s rapid and/or sustained clinical antidepressant effects [19]. Biotransformation of ketamine yields pharmacologically active metabolite (*2R*,*6R*)-hydroxynorketamine (HNK), which is detected in the brain minutes after systemic ketamine administration [20] and exhibits a preclinical NMDAR inhibition-independent ketamine-like antidepressant profile [21] without dissociative side effects in humans [22]. HNK also rapidly potentiates *in vivo* measures of cortical high-frequency neural activity (i.e., gamma oscillatory activity) in mice [21,23], and is currently being assessed for antidepressant efficacy in a phase II clinical trial.

Emerging evidence indicates the initial glutamate surge may converge with canonical NMDAR activation-dependent plasticity to persistently facilitate therapeutic effects by leveraging metaplastic mechanisms (“priming mechanisms” herein) that prime synapses for subsequent changes in strength [6]. *In vitro* perfusion of ketamine or HNK onto hippocampal slices during electrophysiology recordings is an effective model for differentiating their acute pharmacological action as there is no evidence of ketamine metabolism in brain tissue [20], which can be attributed to a near absence of the cytochrome P450s responsible for the formation of HNK [24]. Using such a perfusion model, we reported that HNK, but not ketamine [25], rapidly enhances glutamate release probability to potentiate excitatory neurotransmission at the hippocampal Schaffer collateral-CA1 (SC-CA1) synapse [21,26,27]. However, the ability to resolve the involvement of distinct mediators of the rapid versus sustained antidepressant synaptic action of ketamine or HNK is limited by inherent features of perfusion paradigms such as experimental stimulation that alters slice activity which could preclude antidepressant-relevant action by blocking spontaneous NMDAR activity proposed to facilitate ketamine’s rapid action [15]. Thus, an *in vitro* model that permits the delineation of temporal-specific mediators of ketamine/HNK’s *in vivo* antidepressant synaptic actions would aid in accelerating the resolution of the debate surrounding ketamine’s antidepressant pharmacology, and could facilitate the discovery of novel druggable mechanisms. Therefore, we developed a slice incubation model with pharmacological fidelity to the synaptic effects of bath-applied and systemically-administered HNK, finding that rapid presynaptic potentiation following HNK exposure persistently primes SC-CA1 plasticity.

## MATERIALS/METHODS

See Supplemental Materials/Methods for complete details.

### Animals

Eight-week-old male/female CD-1 mice (Charles River) were housed in groups of four/cage upon arrival. Animals acclimated to the vivarium (University of Maryland, Baltimore) for at least one week after arrival and were maintained on a 12 hr light/dark cycle. Food/water were provided *ad libitum*. All experiments were approved by the University of Maryland Baltimore IACUC and were completed per the latest NIH Guide for the Care and Use of Laboratory Animals.

### Materials

HNK was synthesized and chirality/purity characterized by the National Center for Advancing Translational Sciences, dissolved in 0.9% saline vehicle (VEH), and was either bath-applied to slices, incubated with slices, or administered intraperitoneally in a volume of 7.5 mL/kg of body mass.

### Hippocampal electrophysiology

#### Ex vivo treatment

After one week of acclimation, nine-week-old mice were randomly assigned to one of two treatment groups (VEH or HNK), and at 09:00 animals were treated with VEH or 10 mg/kg HNK by a male experimenter [28]. Two mice/cage received each of the treatments. Animals were anesthetized with isoflurane and euthanized for recordings to begin 3 hr post-treatment.

#### Slice preparation

Hippocampal slice preparation and electrophysiology experiments were conducted as previously described [29,30]. Brains were removed and rapidly submerged in carbogenated (95% O_2_/5% CO_2_), ice-cold dissection artificial cerebrospinal fluid (ACSF). Following 90 min recovery in recording ACSF, slices were transferred to a submersion-type chamber and continuously perfused (1.5 mL/min; Ismatec Reglo ICC Digital Pump; Cole-Parmer) with recording ACSF for recording experiments.

### Experimental design and statistical analyses

Experiments were performed and analyzed by an experimenter blind to treatment groups. All statistical analysis/graphic production were completed using GraphPad Prism version 10.1.2 (GraphPad Software Inc.).

## RESULTS

For full details of statistical outcomes, see **Table S1**.

### (2R,6R)-HNK rapidly potentiates SC-CA1 synaptic efficacy and persistently primes synaptic plasticity

We assessed the impact of bath-applied HNK on synaptic efficacy at the hippocampal SC- CA1 synapse, a synapse well-established to involve depression phenotypes and antidepressant-relevant changes in synaptic strength [5,31]. We used a concentration (10 µM) similar to levels found in the hippocampus minutes after a 10 mg/kg dose [21,32] that rapidly potentiates SC-CA1 synaptic efficacy *in vitro* coincident with an NMDAR inhibition-independent enhanced probability of glutamate release [21,26,27,32]. The exposure length (60-min) is consistent with the duration of a ketamine or HNK infusion [22]. In slices from both male and female mice, baseline paired-pulse ratio (PPR) values were determined, and evoked fEPSPs were monitored for 10-min before slices were exposed to VEH/HNK for 60-min, then VEH/HNK were washed out for the subsequent 35-min. PPR was then reassessed (**Fig. 1A-B**).

**Figure 1.**
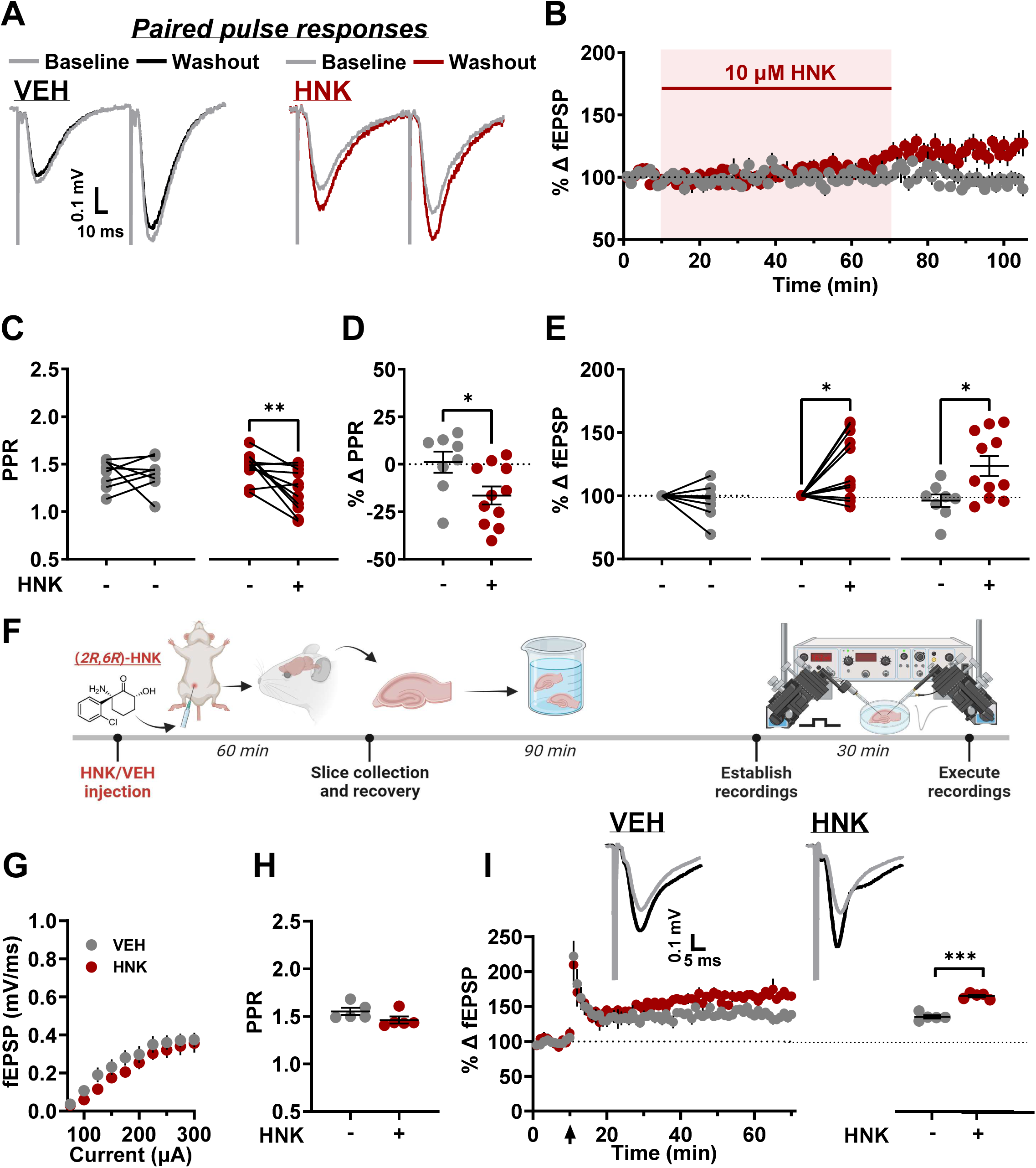
(*2R*,*6R*)-HNK rapidly potentiates synaptic efficacy and persistently primes synaptic metaplasticity. Following a 10 min baseline, hippocampal slices were continuously perfused with either vehicle (VEH) or (*2R*,*6R*)-hydroxynorketamine (HNK; 10 µM) for 60 min while the field excitatory postsynaptic potential (fEPSP) recorded from the Schaffer collateral-CA1 (SC-CA1) synapse was monitored followed by a 35 min washout. PPR was recorded before and after bath application of VEH or HNK. (**A**) Paired-pulse representative traces from baseline (gray) and following bath application of either vehicle (black) or HNK (red). (**B**) Bath application of HNK resulted in synaptic potentiation. (**C, D**) HNK exposure decreased the paired-pulse ratio (PPR; 50 ms interpulse interval), indicative of increased glutamate release probability. (**E**) The fEPSP response was enhanced 35 min after HNK exposure. (**F**) Mice received an intraperitoneal injection of vehicle or 10 mg/kg HNK and were euthanized for slice electrophysiology recordings to begin 3 hr after treatment. (**G**) HNK had no effect on basal synaptic transmission 3 hr after administration as revealed by the unaltered I/O relationship. (**H**) HNK did not alter PPR 3 hr after administration. (**I**) LTP was more readily formed in slices collected from mice 3 hr after HNK administration compared to VEH-treated mice. Arrow at t=10 min denotes 4×100 Hz high-frequency stimulation. Traces are composed of representative sweeps from 5 min pre-tetanus (grey) and 56–60 min post-tetanus (black). Data are the mean ± SEM. * *p*<0.05; ** p<0.01; *** *p*<0.001 as indicated by Holm-Šídák *post-hoc* comparisons. See Table S1 for complete details on the statistical analyses and precise group sizes.

*Post-hoc* analysis revealed that HNK exposure significantly reduced the PPR following wash-out (**Fig. 1C-D**), indicating HNK enhanced glutamate release probability. This indicator of an increase in glutamate release probability was associated with potentiated synaptic transmission following wash-out (**Fig. 1E**; regression model: F_(2,16)_=3.901, p=0.042, R^2^ value=0.33; HNK treatment effect: F_(1,16)_=7.690, p=0.014). We note a slower onset of HNK potentiation than we previously reported in hippocampal slices collected from C57BL/6J mice or Sprague Dawley/Wistar rats [21,26,28,33], which we attribute to the use of CD1 mice. No significant effect of sex was observed in wash-in experiments (Main effect of HNK: F_(1,15)_ = 7.447, p=0.016; sex main effect: F_(1,15)_ = 0.2225, p=0.64; HNK x sex interaction: F_(1,15)_ =1.089, p=0.31). Therefore, experiments described herein were conducted with slices collected from female mice.

We recently reported that a 10 mg/kg dose of (*R*,*S*)-ketamine leverages NMDAR activation-dependent plasticity mechanisms to prime SC-CA1 to persistently lower the threshold for LTP [30]. Thus, we assessed if HNK also activates synaptic priming mechanisms by dosing animals with VEH or 10 mg/kg HNK and assessing LTP 3 hr later (**Fig. 1F**). Basal synaptic transmission and PPR were unaffected 3 hr after the HNK dose (**Fig. 1G-H**). However, LTP was significantly greater in slices collected from HNK-treated animals than LTP from slices of VEH-treated mice (**Fig. 1I**), consistent with HNK activating synaptic priming mechanisms to lower the threshold for LTP.

### Hippocampal slice incubation recapitulates the rapid and sustained synaptic actions of bath-applied and systemically administered (2R,6R)-HNK

To develop an *in vitro* model that allows delineation of the mechanisms leveraged by HNK to rapidly potentiate synaptic strength from those involved in sustained priming, we tested if enhanced synaptic transmission and decreased PPR observed 35-min after HNK exposure are still observed when slices are incubated with HNK in the absence of electrical stimulation. Specifically, slices were incubated with HNK for 1 hr in a holding chamber, and subsequently washed with HNK-free ACSF for 35-min while baseline responses were established in a recording chamber (**Fig. 2A**). Following washout, the rapid effects observed when slices were exposed in the recording chamber were recapitulated where HNK-incubated slices exhibited enhanced basal synaptic transmission compared to VEH-incubated slices. (**Fig. 2B**), reduced PPR (**Fig. 2C**), and, interestingly, reduced LTP (**Fig. 2D**). Multiple linear regression analyses revealed that the reduction in PPR was associated with the increase in maximal fEPSP elicited in the input/output (I/O) curve (regression model: F_(2,16)_=6.553, p=0.0083, R^2^ value=0.45; HNK treatment effect: F_(1,16)_=13.10, p=0.0023; predictor variable PPR effect: F_(1,16)_=4.605, p=0.048).

**Figure 2.**
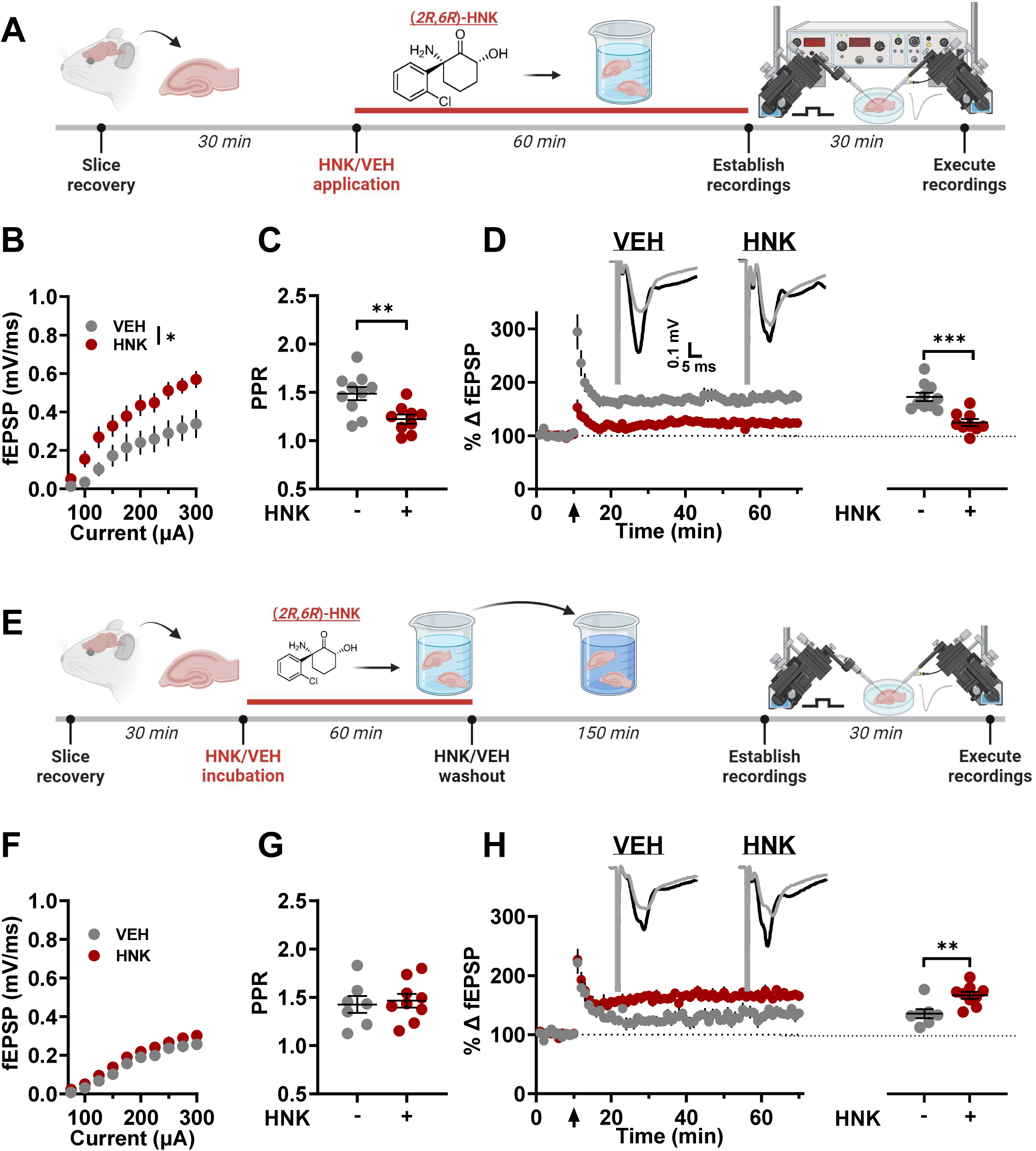
Hippocampal slice incubation recapitulates the rapid and sustained synaptic actions of bath-applied and systemically administered (*2R*,*6R*)-HNK. (**A**) Following 30 min recovery in an ACSF-containing, drug-free holding chamber, hippocampal slices were treated with either vehicle (VEH) or (*2R*,*6R*)-hydroxynorketamine (HNK; 10 µM) for 60 min followed by a 35 min washout while the field excitatory postsynaptic potential (fEPSP) recorded from the Schaffer collateral-CA1 (SC-CA1) synapse was established in the recording chamber. (**B**) HNK potentiated the basal fEPSP response as indicated by increased input/output curve response and (**C**) increased glutamate release probability as indicated by reduced paired-pulse ratio (PPR; 50 ms interpulse interval). (**D**) The capacity to form long-term potentiation (LTP) was reduced following 35 min wash-out. (**E**) Following 30 min recovery in an ACSF-containing, drug-free holding chamber, hippocampal slices were treated with either VEH or HNK (10 µM) for 60 min followed by a 150 min washout in a separate holding chamber before a 30 min period where fEPSP responses from the SC-CA1 synapse were established in the recording chamber (3 hr total HNK washout). (**F**) HNK did not affect basal synaptic transmission 3 hr after exposure as revealed by the unaltered I/O relationship. (**G**) PPR was unchanged 3 hr after HNK exposure. (**H**) LTP was more readily formed in slices exposed to HNK 3 hr earlier compared to VEH- incubated slices. Arrow at t=10 min denotes 4×100 Hz high-frequency stimulation. Traces are composed of representative sweeps from 5 min pre-tetanus (grey) and 56–60 min post-tetanus (black) from treatment groups. Vertical calibration bar (0.1 mV) and horizontal calibration bar (5 ms) are consistent for all traces. Data are the mean ± SEM. * *p*<0.05; ** p<0.01; *** *p*<0.001 as indicated by Holm-Šídák *post-hoc* comparisons except in **B** in which * denotes the main effect of HNK. See Table S1 for complete details on the statistical analyses and precise group sizes.

We also assessed the sustained priming effect of HNK observed on *ex vivo* LTP 3 hr after administration by incubating slices with HNK for 1 hr followed by 3 hr washout in a drug-free chamber (**Fig. 2E**). These wash-in/wash-out periods are in line with the *in vivo* pharmacokinetics of HNK in CD-1 mice where brain exposure to the drug is limited to approximately 1 hr post-intraperitoneal administration [34]. Consistent with our *ex vivo* experiment, 3 hr after HNK incubation basal synaptic transmission (**Fig. 2F**) and PPR (**Fig. 2G**) were unchanged, but HNK- incubated slices exhibited significantly greater LTP than VEH-incubated slices (**Fig. 2H**). While LTP was still observed in VEH-incubated slices, we note a reduction of LTP magnitude in slices 3 hr after VEH exposure compared to 35-min after, an effect likely due to the duration of the experiment before completion of our assessments (∼6-7 hr post-euthanization). However, baseline transmission and PPR in VEH-incubated slices were not significantly different between the time points (I/O: main effect of time point: F_(1,15)_=0.6000, p=0.45; PPR: t_15_=0.57, p=0.58).

### Rapid synaptic potentiation evoked by (2R,6R)-HNK occurs independently of NMDAR activation, but persistent synaptic priming is NMDAR activation-dependent

We have previously shown that HNK, unlike ketamine, rapidly potentiates synaptic transmission through an NMDAR inhibition-independent mechanism [21,26,32], but the involvement of the NMDAR in its sustained action remains untested. Thus, we preincubated slices with an NMDAR antagonist to assess its time-sensitive pharmacology. Specifically, after 20 min recovery, we pretreated slices with VEH or 50 µM D-APV for 10-min before exposure to VEH/HNK (**Fig. 3A**). Slices were then coincubated with D-APV for the subsequent 60-min before establishing baseline responses in a 35-min wash-out period. Consistent with previous studies [21,26], D-APV pretreatment did not prevent HNK’s ability to enhance basal synaptic transmission (**Fig. 3B**) or affect HNK’s ability to reduce PPR (**Fig. 3C**). Likewise, the reduction in LTP after HNK exposure was unaffected by D-APV pretreatment (**Fig. 3D**).

**Figure 3.**
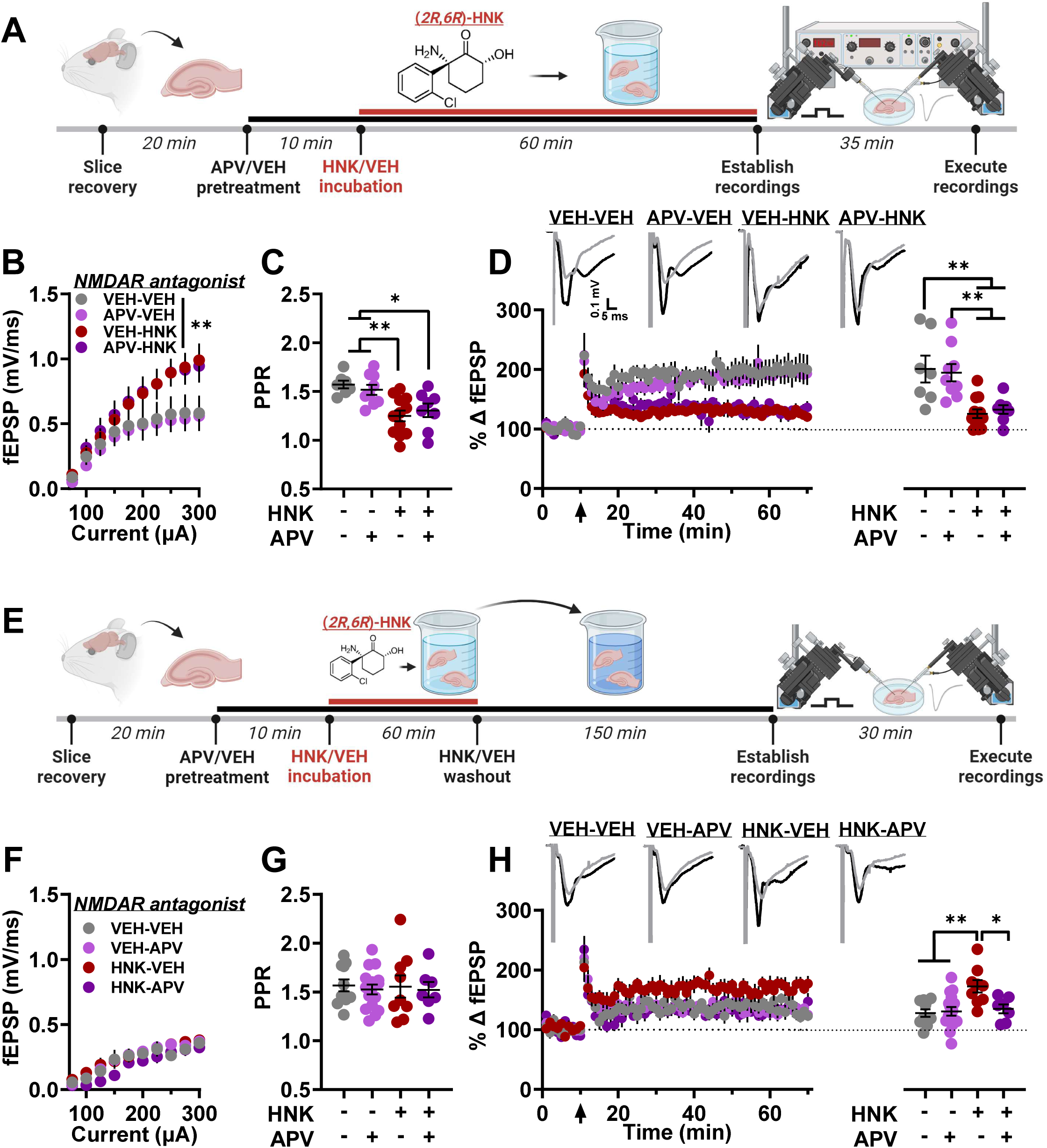
Rapid potentiation of synaptic efficacy by (*2R*,*6R*)-HNK occurs independently of NMDAR activation, but persistent synaptic priming is NMDAR activation-dependent. (**A**) Following 20 min recovery in an ACSF-containing, drug-free holding chamber, hippocampal slices were treated with either vehicle (VEH) or NMDAR antagonist D-APV (50 µM) for 10 min followed by VEH or (*2R*,*6R*)-hydroxynorketamine (HNK; 10 µM) treatment at t=30 min. Slices were coincubated for 60 min followed by a 35 min washout while the field excitatory postsynaptic potential (fEPSP) recorded from the Schaffer collateral-CA1 (SC-CA1) synapse was established. (**B**) HNK potentiated the basal fEPSP response as indicated by increased input/output curve response and (**C**) increased glutamate release probability as indicated by reduced paired-pulse ratio (PPR; 50 ms interpulse interval) independent of D-APV pretreatment. (**D**) The capacity to form long-term potentiation (LTP) was reduced following 35 min wash-out independent of D-APV pretreatment. (**E**) Following 20 min recovery in an ACSF-containing, drug-free holding chamber, hippocampal slices were treated with either VEH or NMDAR antagonist D-APV (50 µM) for 10 min followed by treatment with VEH or HNK (10 µM) at t=30 min. Slices were coincubated for 60 min followed by a 150 min HNK washout in a separate holding chamber before a 30 min period where fEPSP responses from the SC-CA1 synapse were established in the recording chamber (3 hr total HNK washout). (**F**) HNK/D-APV did not affect basal synaptic transmission 3 hr after exposure as revealed by the unaltered I/O relationship. (**G**) PPR was unchanged by HNK/D-APV. (**H**) LTP was more readily formed in slices exposed to HNK 3 hr earlier compared to VEH-incubated slices, and this effect was blocked by posttreatment, but not pretreatment with D-APV. Arrow at t=10 min denotes 4×100 Hz high-frequency stimulation. Traces are composed of representative sweeps from 5 min pre-tetanus (grey) and 56–60 min post-tetanus (black) from treatment groups. Vertical calibration bar (0.1 mV) and horizontal calibration bar (5 ms) are consistent for all traces. Data are the mean ± SEM. * *p*<0.05; ** p<0.01; *** *p*<0.001 as indicated by Holm-Šídák *post-hoc* comparisons except in **B** in which ** denotes the main effect of HNK. See Table S1 for complete details on the statistical analyses and precise group sizes.

We next asked if NMDAR activity was required for the sustained synaptic priming evoked by HNK by incubating a subset of slices with D-APV for 10-min before 1 hr VEH/HNK coexposure in a holding chamber followed by a 3 hr D-APV and HNK washout in a second drug-free ACSF holding chamber (treatment groups: VEH-VEH or HNK-VEH). Another subset was continuously incubated with D-APV-containing, HNK-free ACSF for 150 min in the second holding chamber (treatment groups: VEH-APV or HNK-APV; **Fig. 3E**). No treatment groups exhibited significant changes in I/O (**Fig. 3F)** or PPR (**Fig. 3G**), but slices that received a 3 hr HNK wash-out in D-APV-free ACSF exhibited enhanced LTP, an effect not observed in slices which were exposed to D-APV after HNK incubation (**Fig. 3H**). Overall, these results are consistent with HNK rapidly potentiating SC-CA1 neurotransmission in an NMDAR-independent manner to facilitate synaptic signaling that recruits NMDAR activation-dependent synaptic priming mechanisms that persistently lower the threshold for LTP.

### Rapid potentiation of synaptic efficacy by (2R,6R)-HNK requires the activity of adenylyl cyclase isoform 1, but not isoform 5

We recently reported that HNK leverages AC-cAMP-PKA-dependent signaling to rapidly potentiate synaptic transmission at the SC-CA1 synapse by increasing glutamate release probability [33], a finding that is in line with the established role of AC-cAMP-PKA-dependent signaling in hippocampal glutamate release probability [35,36]. Accordingly, we used the incubation protocol to assess if our model can also detect HNK’s capacity to engage AC-cAMP- PKA-dependent signaling to impact synaptic activity, and, if so, reveal novel druggable mechanisms. Slices were pretreated with VEH or a pan-AC inhibitor, 10 µM SQ22536, for 10- min and were then exposed to VEH/HNK for 60-min, followed by a 35-min drug washout in the recording chamber **(Fig. 4A)**. Incubation of slices with SQ22536 before HNK blocked rapid enhancement of basal synaptic transmission (**Fig. 4B**) and reduced PPR (**Fig. 4C**). In a separate experiment, the decrease in LTP 35-min after HNK exposure was blocked by SQ22536 pretreatment (**Fig. 4D**). These data show that our incubation model can delineate the AC-cAMP- PKA signaling pathway leveraged by HNK to promote its acute synaptic effects at the SC-CA1 synapse.

**Figure 4.**
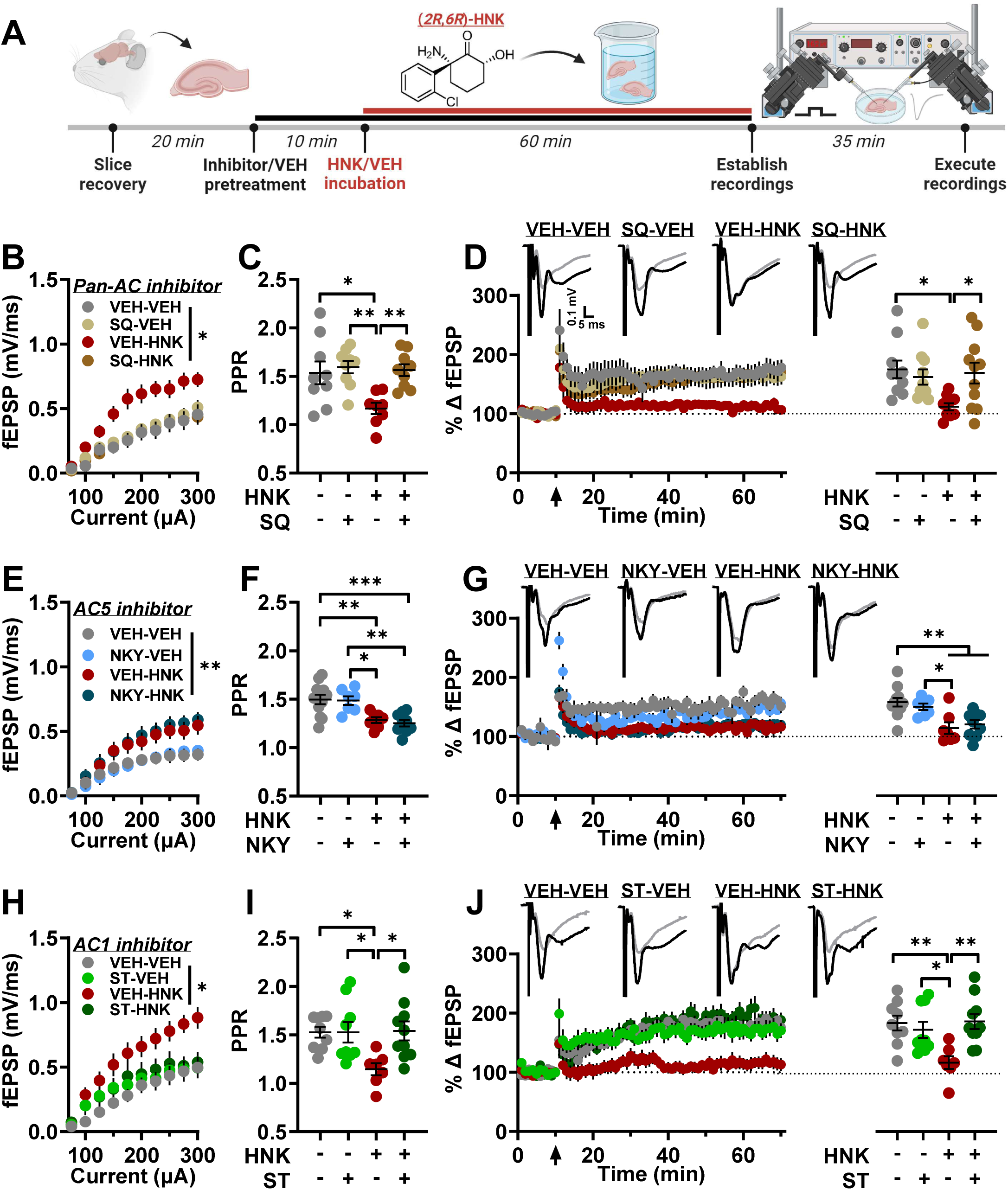
Rapid potentiation of synaptic efficacy by (*2R*,*6R*)-HNK requires the activity of adenylyl cyclase isoform 1, but not isoform 5. (**A**) Following 20 min recovery in an ACSF-containing, drug-free holding chamber, hippocampal slices were treated with either vehicle (VEH) or either a non-competitive pan-adenylyl cyclase (AC) inhibitor (SQ22536; 10 µM; **B-D**), selective AC5 inhibitor (NKY80; 10 µM; **E-G**), or selective AC1 inhibitor (ST034370; 1 µM; **H-J)** for 10 min followed by VEH or (*2R*,*6R*)-hydroxynorketamine (HNK; 10 µM) treatment at t=30 min. Slices were coincubated for 60 min followed by a 35 min washout while the field excitatory postsynaptic potential (fEPSP) recorded from the Schaffer collateral-CA1 (SC-CA1) synapse was established. (**B**) HNK potentiated the basal fEPSP response as indicated by increased input/output curve response and (**C**) increased glutamate release probability as indicated by reduced paired-pulse ratio (PPR; 50 ms interpulse interval) in a SQ22536-dependent manner. (**D**) The capacity to form long-term potentiation (LTP) was reduced following 35 min wash-out in an SQ22536-dependent manner. (**E**) HNK potentiated the basal fEPSP response as indicated by increased input/output curve response and (**F**) increased glutamate release probability as indicated by reduced PPR in an AC5-independent manner. (**G**) The capacity to form LTP was reduced following 35 min wash-out in an AC5-independent manner. (**H**) HNK potentiated the basal fEPSP response as indicated by increased input/output curve response and (**I**) increased glutamate release probability as indicated by reduced PPR in an AC1-dependent manner. (**J**) The capacity to form LTP was reduced following 35 min wash-out in an AC1- dependent manner. Arrow at t=10 min denotes 4×100 Hz high-frequency stimulation. Traces are composed of representative sweeps from 5 min pre-tetanus (grey) and 56–60 min post-tetanus (black) from treatment groups. Vertical calibration bar (0.1 mV) and horizontal calibration bar (5 ms) are consistent for all traces. Data are the mean ± SEM. * *p*<0.05; ** p<0.01; *** *p*<0.001 as indicated by Holm-Šídák *post-hoc* comparisons except in **B**, **E**, **H** in which * or ** denotes the main effect of HNK. See Table S1 for complete details on the statistical analyses and precise group sizes. Abbreviations: NKY, NKY80; SQ, SQ22536; ST, ST034307.

Group III AC isoform AC5 and group I AC isoform AC1 [37] play critical roles in synaptic plasticity in CA3/CA1 [38] and are linked to neuropsychiatric conditions [39]. Thus, we pretreated slices with VEH or the AC5 selective inhibitor, NKY80 (10 µM), for 10-min before incubation in VEH/HNK for 1 hr. Following washout, NKY80 pretreatment did not prevent enhanced basal synaptic transmission in slices co-incubated with HNK (**Fig. 4E**), reduced PPR (**Fig. 4F**), or LTP magnitude (**Fig. 4G**), indicating that AC5 activity is not required for the observed rapid synaptic effects of HNK at SC-CA1.

We next pretreated slices with VEH or 1 µM ST034307, a selective inhibitor of Ca^2+^/calmodulin-activated AC1 [40], followed by coincubation with either VEH/HNK for 1 hr to probe the necessity of the AC1 isoform in the rapid effects of HNK. Consistent with our SQ22536 experiment, ST034307 preincubation completely prevented the enhanced I/O curve 35-min after HNK washout (**Fig. 4H**), the PPR reduction (**Fig. 4I**), and LTP decrease (**Fig. 4J**). Thus, these findings are consistent with HNK leveraging AC1-dependent signaling to facilitate rapid potentiation of neurotransmission at SC-CA1.

### (2R,6R)-HNK leverages protein kinase A activity-dependent signaling to rapidly potentiate synaptic efficacy, and this potentiation facilitates persistent synaptic plasticity priming

Next, we assessed the ability of our incubation model to directly distinguish the temporally progressive mechanisms that promote HNK’s rapid and sustained antidepressant-relevant synaptic plasticity. First, slices were preincubated for 10-min with VEH or 10 µM H-89, a cell-permeable inhibitor of PKA, before 1 hr VEH/HNK exposure, followed by a 35-min washout (**Fig. 5A**). Consistent with our findings in which H-89 preincubation blocked the potentiating effects of bath-applied HNK [33], we found that slices incubated with H-89 before HNK exposure did not exhibit enhanced I/O responses (**Fig. 5B**), decreased PPR (**Fig. 5C**), or reduced LTP (**Fig. 5D**) following washout.

**Figure 5.**
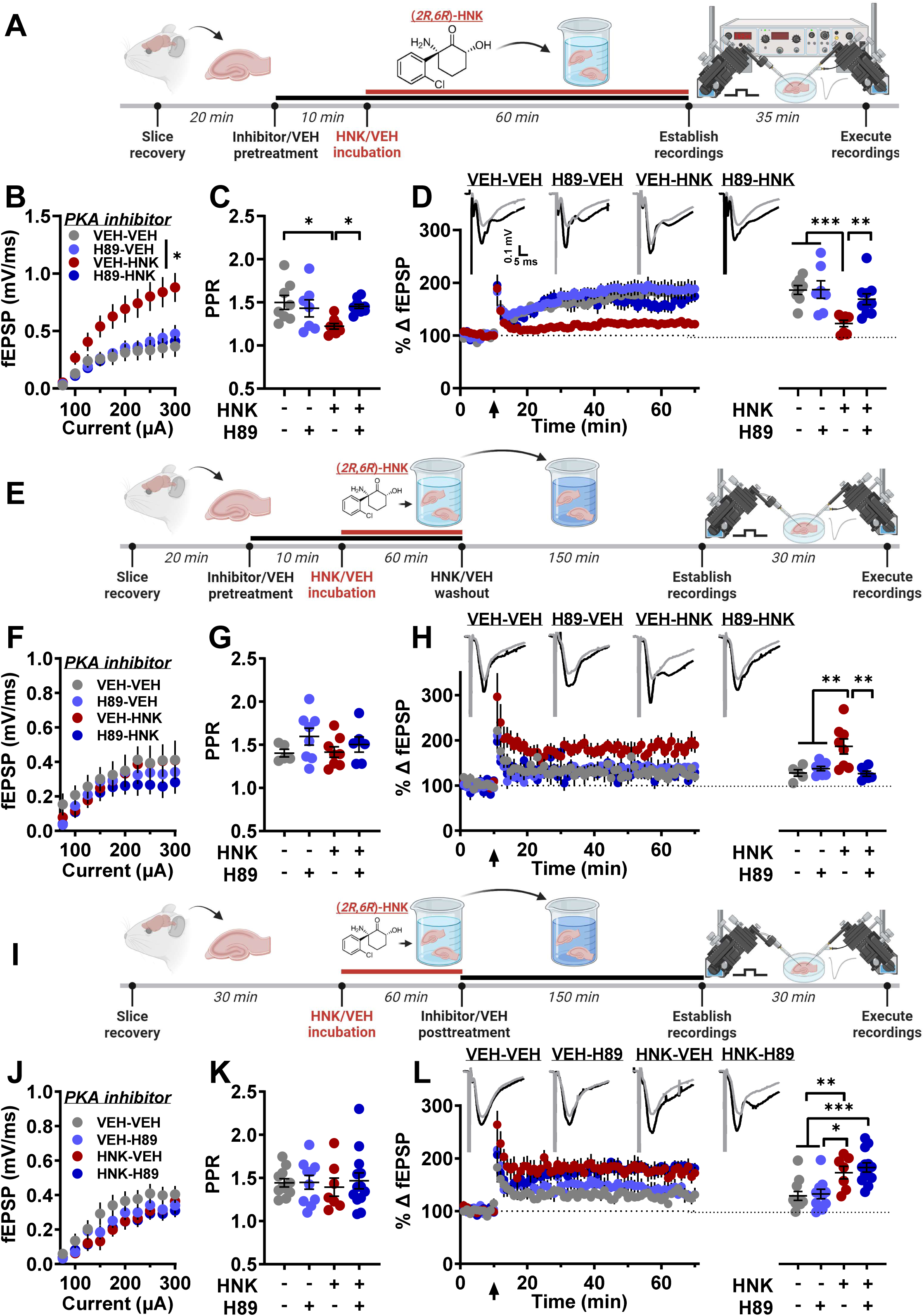
(*2R*,*6R*)-HNK requires protein kinase A activity-dependent signaling to rapidly potentiate synaptic efficacy, and this potentiation facilitates persistent synaptic plasticity priming. (**A**) Following 20 min recovery in an ACSF-containing, drug-free holding chamber, hippocampal slices were treated with either vehicle (VEH) or protein kinase A (PKA) inhibitor H-89 (10 µM) for 10 min followed by VEH or (*2R*,*6R*)-hydroxynorketamine (HNK; 10 µM) treatment at t=30 min. Slices were coincubated for 60 min followed by a 35 min washout while the field excitatory postsynaptic potential (fEPSP) recorded from the Schaffer collateral-CA1 (SC-CA1) synapse was established. (**B**) HNK potentiated the basal fEPSP response as indicated by increased input/output curve response and (**C**) increased glutamate release probability as indicated by reduced paired-pulse ratio (PPR; 50 ms interpulse interval) in an H-89-dependent manner. (**D**) The capacity to form long-term potentiation (LTP) was reduced following 35 min wash-out in an H-89-dependent manner. (**E**) Following 20 min recovery in an ACSF-containing, drug-free holding chamber, hippocampal slices were treated with either VEH or PKA inhibitor H-89 (10 µM) for 10 min followed by treatment with VEH or HNK (10 µM) at t=30 min. Slices were coincubated for 60 min followed by a 150 min HNK/H-89 washout in a separate holding chamber before a 30 min period where fEPSP responses from the SC-CA1 synapse were established (3 hr total HNK/H-89 washout). (**F**) HNK/H-89 did not affect basal synaptic transmission 3 hr after exposure as revealed by the unaltered I/O relationship. (**G**) PPR was unchanged by HNK/H-89. (**H**) LTP was more readily formed in slices exposed to HNK 3 hr earlier compared to VEH-incubated slices, and this effect was blocked by pretreatment with H-89. (**I**) Following 30 min recovery in an ACSF-containing, drug-free holding chamber, hippocampal slices were exposed to either VEH or HNK (10 µM) for 60 min followed by posttreatment in a separate beaker containing VEH or PKA inhibitor H-89 (10 µM) for 150 min. Slices were then moved to the recording chamber for a 30 min VEH/H-89 washout period where fEPSP responses from the SC-CA1 synapse were established (3 hr total HNK washout). (**F**) HNK/H-89 did not affect basal synaptic transmission 3 hr after exposure as revealed by the unaltered I/O relationship. (**G**) PPR was unchanged by HNK/H-89. (**H**) LTP was more readily formed in slices exposed to HNK 3 hr earlier compared to VEH-incubated slices, and this effect was not blocked by posttreatment with H-89. Arrow at t=10 min denotes 4×100 Hz high-frequency stimulation. Traces are composed of representative sweeps from 5 min pre-tetanus (grey) and 56–60 min post-tetanus (black) from treatment groups. Vertical calibration bar (0.1 mV) and horizontal calibration bar (5 ms) are consistent for all traces. Data are the mean ± SEM. * *p*<0.05; ** p<0.01; *** *p*<0.001 as indicated by Holm-Šídák *post-hoc* comparisons except in **B** in which * denotes the main effect of HNK. See Table S1 for complete details on the statistical analyses and precise group sizes.

To determine if rapid synaptic potentiation was required for the activation of priming mechanisms 3 hr after HNK exposure, we preincubated slices with VEH/H-89 for 10-min and then exposed them to either VEH/HNK for 60-min. Following coincubation, slices were moved to a separate, drug-free holding chamber for a 3 hr washout before beginning evoked recordings (**Fig. 5E**). Basal synaptic transmission (**Fig. 5F**) and PPR (**Fig. 5G**) were unaffected by H-89 and/or HNK 3 hr after exposure. Strikingly, H-89 coincubation completely prevented the priming effect of HNK on LTP (**Fig. 5H**), suggesting that rapid enhancement of glutamatergic transmission at SC-CA1 activates priming mechanisms that persistently promote LTP formation.

Lastly, we probed the temporal-specific contribution of PKA activity to the HNK-evoked priming of plasticity. To test this, slices recovered for 30-min and were then incubated with either VEH/HNK for 1 hr (treatment groups: VEH-VEH or HNK-VEH), and then were moved to a separate holding chamber containing either VEH or H-89 (treatment groups: VEH-H89 or HNK- H89) in which they were incubated before evoked recordings began 3 hr after HNK exposure (**Fig. 5I**). I/O (**Fig. 5J**) and PPR (**Fig. 5K**) did not significantly differ in any treatment groups 3 hr after VEH/HNK exposure. Strikingly, H-89 incubation after HNK exposure did not prevent the sustained activation of priming mechanisms as revealed by both HNK-VEH and HNK-H-89 treatment groups exhibiting significantly enhanced LTP (**Fig. 5L**). These results underscore the utility of our incubation model to delineate the synaptic machinery engaged in rapid versus sustained synaptic action of rapid-acting antidepressants by revealing that rapid HNK-induced potentiation persistently primes NMDAR activation-dependent plasticity.

## DISCUSSION

We developed a slice incubation model with fidelity to HNK’s rapid effects observed during bath application [21,26,27,33] and its sustained *ex vivo* effects [41] to delineate mechanisms engaged by its time-sensitive antidepressant pharmacology. We used this model to reveal that PKA activity-dependent potentiation of neurotransmission shortly after HNK exposure was required for sustained plasticity priming. We conclude that approaches such as our incubation model can be leveraged to reveal novel druggable mechanisms that engage temporal-sensitive signaling cascades mediating the progression from acute to sustained therapeutic effects of rapid-acting antidepressants.

Impaired synaptic plasticity and transmission have been posited as critical contributors to depression symptomology [31,42,43]. It is largely agreed that plasticity evoked by an antidepressant dose of ketamine rapidly and persistently ameliorates deficits in synaptic transmission, yet the mechanisms by which this plasticity occurs remain highly contested [5,7]. Drug perfusion to brain slices during electrophysiology recordings has been a conventional approach to delineate ketamine’s rapid antidepressant pharmacology. Some studies have found that perfusion of ketamine to hippocampal slices potentiates SC-CA1 synaptic transmission in the absence of electrically evoked activity [44,45]; however, other reports found no acute effects of bath-applied ketamine at this synapse in the absence [46] or presence [47,48] of electrical stimulation. We recently reported that bath-applied ketamine did not alter SC-CA1 synaptic efficacy, but a ketamine veterinary formulation commonly used in bath-application studies did potentiate synaptic transmission, an effect we found was driven by the presence of a preservative (benzethonium chloride) [25,49].

We have reported that HNK application to hippocampal slices potentiates SC-CA1 synaptic efficacy [EC50=3.3 µM; 26,27,33]. Here, we show that potentiated SC-CA1 synaptic strength following bath-applied 10 µM HNK is maintained following 35-min wash-out as revealed by enhancement of the fEPSP and a corresponding augmentation of the I/O curve. Notably, the HNK-elicited synaptic potentiation was correlated to a PPR decrease. Thus, this fEPSP increase is indicative of enhanced glutamate release probability from excitatory SC projections, a finding in line with our reports of HNK decreasing electrically- and optically-evoked SC-CA1 PPR, and increasing miniature (mEPSC) frequency, but not amplitude, recorded from CA1 principal cells during perfusion [26,27,33].

The use of a conventional slice electrophysiology approach (i.e, monitoring responses during drug perfusion) to elucidate the precise synaptic machinery leveraged by ketamine/ketamine metabolites that promote its rapid (i.e., <1 hr) and sustained (i.e., hours-days) therapeutic action is limited by inherent requirements such as stimulation-evoked preclusion of putative ketamine’s rapid antidepressant pharmacology [15] or experimental-induced slice decay occurring after hours of monitoring responses. We endeavored to develop a model that can distinguish the time-sensitive synaptic action of rapid-acting antidepressants. Consistent with our bath-application experiments, slices incubated with 10 µM HNK in the absence of stimulation exhibited enhanced I/O responses and reduced PPR following a 35-min washout. The maximal I/O response was correlated to the PPR decrease after wash-out. We recently reported that HNK’s rapid bath-applied effects at SC-CA1 were prevented by pretreatment with cell-permeable pan-inhibitors of AC (SQ22536) or PKA (H-89) activity [33]. We verified the pharmacological validity of our incubation model by showing that the HNK-associated changes in synaptic activity following 35-min washout were also blocked by pretreatment with these inhibitors. Notably, we extend our previous results by finding that AC1 activity, but not AC5, is required for the rapid effects of HNK. AC1 is highly expressed in CA1/3 subfields [38] and functions as the primary Ca^2+^ stimulation-dependent AC contributor to cAMP accumulation in CA3 [50], consistent with HNK leveraging the trigger for vesicular fusion, Ca^2+^, to enhance glutamate release probability and increase CA3-CA1 synaptic efficacy. Additionally, postsynaptic AC1 is critical for synaptic plasticity as studies found AC1 knockout impaired hippocampal-dependent memory and *ex vivo* SC-CA1 LTP [51] whereas AC1 overexpression enhanced hippocampal-dependent learning and SC-CA1 LTP recorded *in vivo* [52]. These findings support the earlier findings of NMDAR- independent, cAMP-dependent antidepressant actions of HNK [53]. Thus, the physiological function of AC1 in synaptic physiology, combined with our results here, supports the AC1 isoform as an attractive target in neuropsychiatric drug development [54].

Little has been reported on the acute, *in vitro* effects of HNK on the subsequent ability to form LTP. One study found reduced SC-CA1 LTP when slices were pretreated with 10 µM HNK for 2 hr followed by continuous HNK perfusion during recording, an effect attributed to direct inhibitory action of HNK on the NMDAR [55]. This interpretation is consistent with the induction of LTP at SC-CA1 being NMDAR-dependent [56,57]. Yet studies indicate that HNK’s binding affinity for the NMDAR at 10 µM as measured by MK-801 displacement is negligible (K_i_ > 100 µM; [21,58,59]). Functional effects of HNK on NMDAR-related responses occur at concentrations much greater than those required for antidepressant-like effects (e.g., HNK IC_50_=211.9 µM for SC-CA1 NMDAR-mediated fEPSP; IC_50_=63.7 µM for CA1 principal cell mEPSC amplitude [21,32]; NMDAR open channel blocker at 50 µM but not 10 µM in hippocampal neuron culture [60]). Further, we reported that HNK augments SC-CA1 synaptic efficacy in the presence of NMDAR inhibitor D-APV [26]. Thus, the potentiating effects induced by HNK at an antidepressant-relevant concentration likely arise independently of the NMDAR. Here we report decreased LTP 35-min after HNK wash-out while indicators of presynaptic potentiation persist, even when slices were coincubated with an NMDAR antagonist. This LTP reduction may thus result from elevated synaptic glutamate at the time of LTP induction as it has been reported that elevated synaptic glutamate levels during tetanus can impair LTP via an L-type channel-dependent mechanism [61]. Notably, HNK’s action to rapidly enhance *in vitro* brain-derived neurotropic factor release requires L-type channel activity [62]. Therefore, while our results are consistent with a presynaptic locus for the initiation of HNK’s effects, the reduction in LTP suggests a time-sensitive progression to a postsynaptic site of action shortly after exposure (≤35-min), which warrants future investigation.

The timing, intensity, and/or frequency of stimulus events can be exploited to prime synapses to respond distinctly upon plasticity induction hours-days after primer exposure (reviewed in detail in [6]), resulting in a continuous shift in the propensity for plasticity toward a state that either favors synaptic potentiation or depression [63] without impacting evoked basal activity [64,65] via a phenomenon collectively described as metaplasticity [66]. The NMDAR is a critical cellular regulator of primer-activated metaplasticity mechanisms. For instance, the threshold for CA1 homosynaptic LTP is enhanced 60-90 min after NMDAR activation [67], whereas NMDAR blockade with low ketamine concentrations can subsequently promote LTP [46,68]. Thus, the capacity for a subanesthetic ketamine dose to lower the threshold for SC-CA1 LTP *in vivo* [41] and *ex vivo* [30,47,69–71] 3-24 hr post-treatment may be due to its inhibitory action on the NMDAR subsequently shifting the synapse towards a state that favors LTP. This is consistent with our recent report showing the priming effect of ketamine on *ex vivo* SC-CA1 LTP measured 24 hr after treatment required NMDAR activity at the time of drug administration [30]. However, ketamine metabolite HNK has also been demonstrated to prime the SC-CA1 synapse for LTP when recorded *in vivo* 3.5 hr later [41].

Here, we similarly found *ex vivo* LTP was more readily formed 3 hr after *in vivo* HNK treatment. We recapitulated this finding in our incubation model, showing that synaptic priming was dependent on NMDAR activity only after HNK treatment. To the best of our knowledge, this is the first report of an *in vitro* incubation model of NMDAR-mediated synaptic priming that exhibits fidelity to *ex vivo* priming. The precise mechanism by which HNK primes plasticity remains unclear, but we used our model to reveal that pretreatment, but not posttreatment, with a PKA inhibitor prevented HNK’s rapid effects, and when these were blocked, LTP priming was not observed. Thus, HNK’s ability to enhance glutamate release probability by leveraging AC1- cAMP-PKA-dependent activity converges with NMDAR activation-dependent priming mechanisms, shifting the basal state of SC-CA1 to persistently favor LTP. Our results here, and elsewhere [30], hence indicate that a focus on psychiatric therapies designed to persistently reduce NMDAR activity may be counterproductive as it appears that numerous rapid-acting antidepressants leverage NMDAR activation-dependent plasticity mechanisms [6,30]. Thus, the exploration of synaptic mechanisms responsible for rapid/sustained antidepressant effects of ketamine and next-generation therapies by using approaches such as our incubation model will facilitate the discovery of novel druggable mechanisms and optimize our ability to exploit them.

## Acknowledgments

We thank Patrick J. Morris and Craig J. Thomas (Division of Preclinical Innovation, National Center for Advancing Translational Sciences, NIH, Rockville, MD) for synthesizing and providing HNK.

## Author contributions

KAB and TDG conceptualized the experiments conducted in the study. KAB and MIA completed the experiments and analyzed the data. KAB wrote the manuscript. All authors critically reviewed the manuscript and approved its final form.

---

**Table S1.**
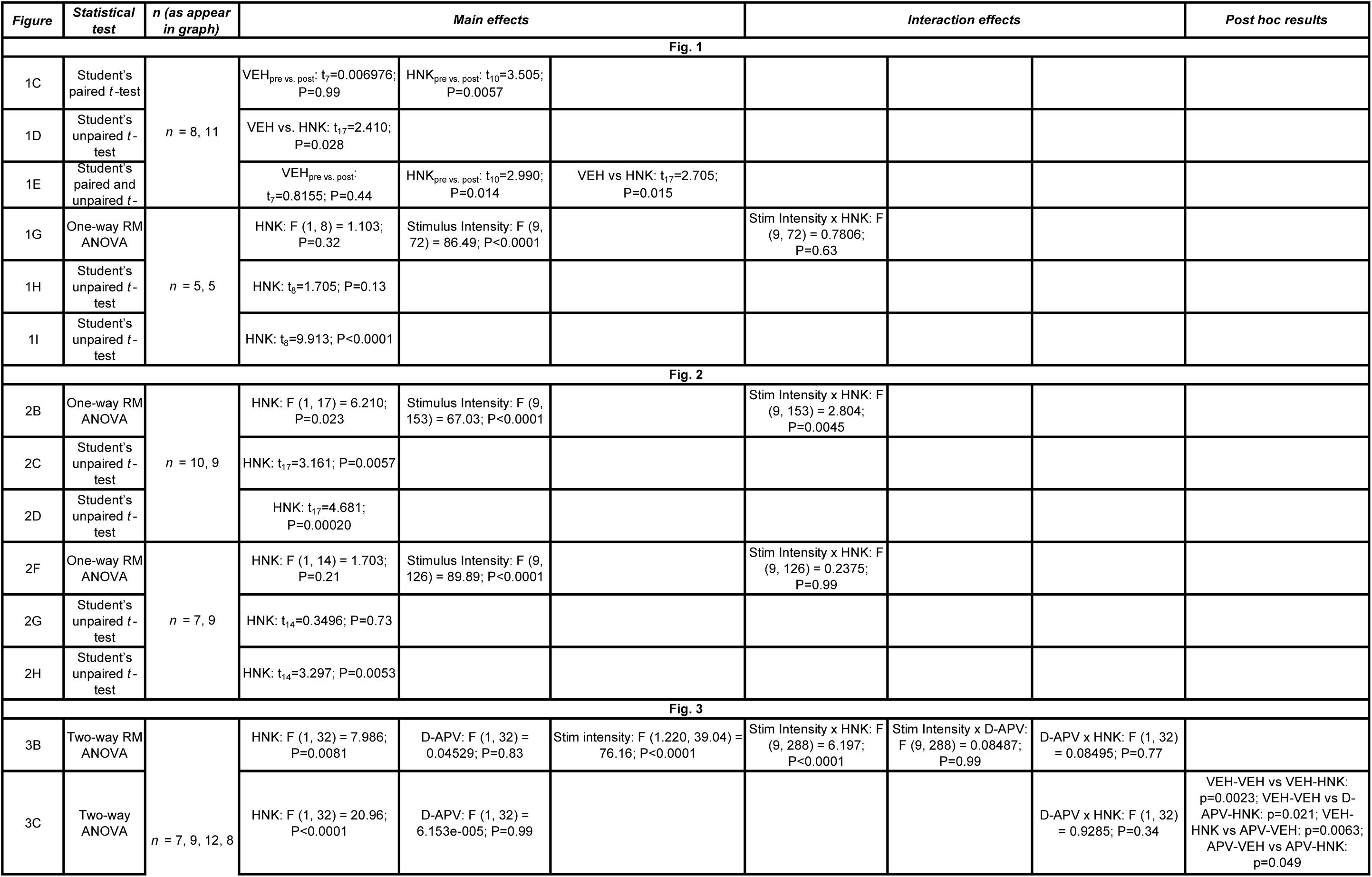
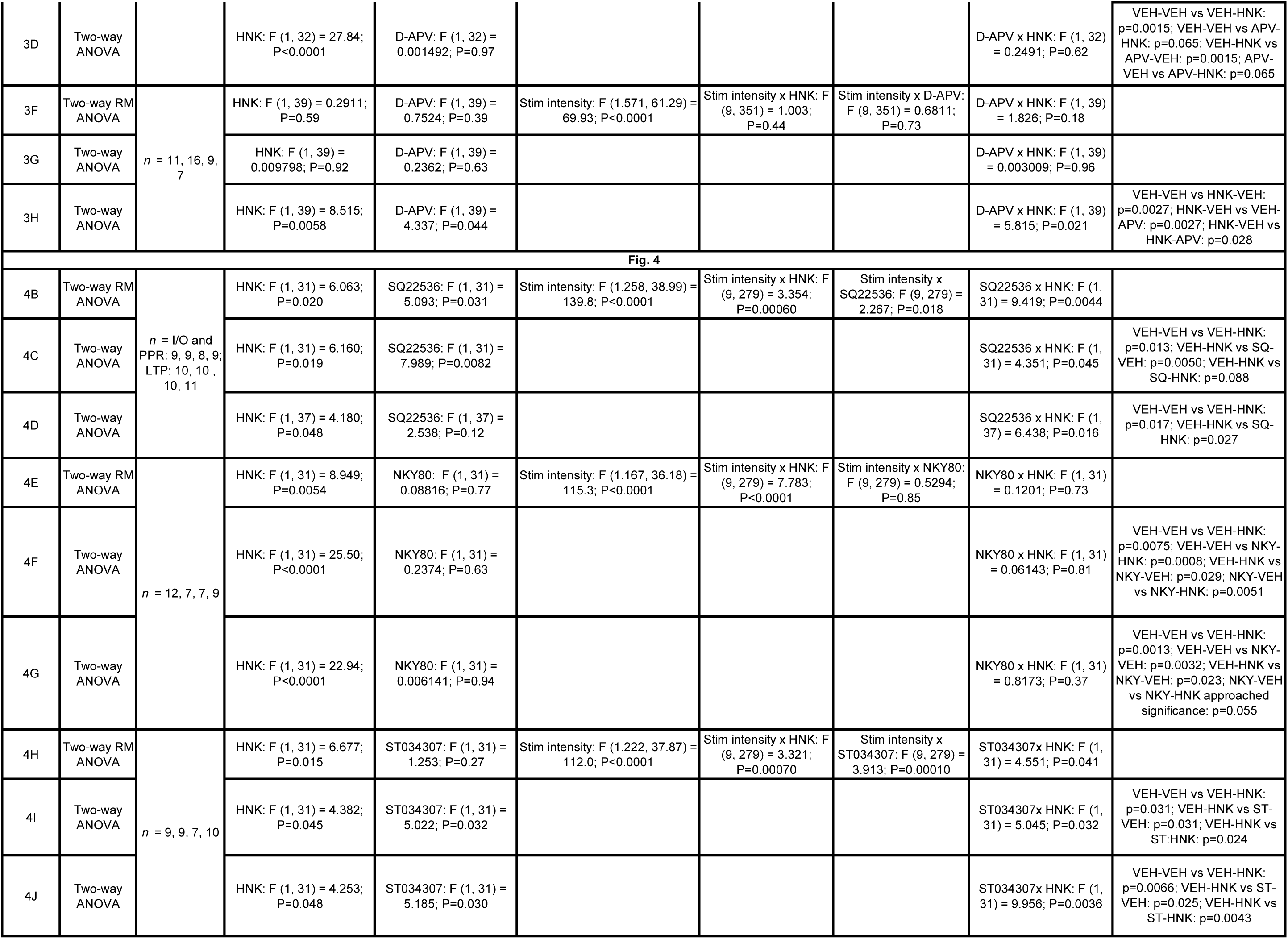
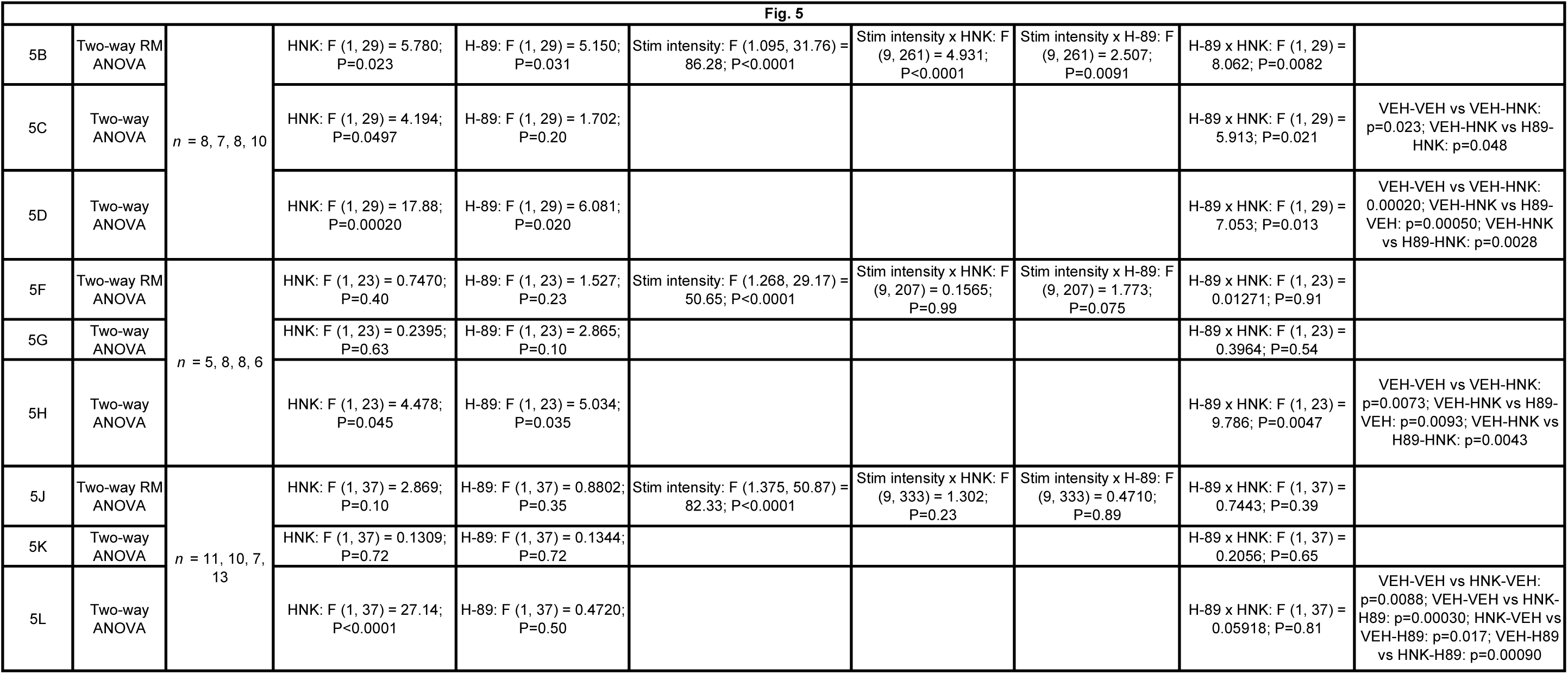
Details on the statistical analyses performed, number of independent observation, main effects, and interaction effects for described results.

## SUPPLEMENTARY MATERIALS AND METHODS

### Materials

H-89 dihydrochloride (Cat. No.: HY-15979A; dissolved in DMSO as VEH), NKY80 (Cat. No: HY-103195; dissolved in DMSO as VEH), ST034307 (Cat. No.: HY-101279; dissolved in DMSO as VEH), and SQ22536 (Cat. No.: HY-100396; dissolved in DMSO as VEH) were purchased from MedChemExpress. DMSO concentrations did not exceed 0.1% in incubation experiments. D-APV (Cat. No. 0106; dissolved in 0.9% saline as VEH) was obtained from Tocris Bioscience. All additional chemicals and reagents used in this study, unless otherwise noted, were of analytical or higher grade and obtained from Sigma-Aldrich.

### Hippocampal slice electrophysiology

After brains were removed from mice and placed in dissection ACSF (120mM NaCl, 3mM KCl, 4mM MgCl_2_, 1mM NaH_2_PO_4_, 26mM NaHCO_3_, and 10mM glucose), the brain was mounted on its dorsal surface and sectioned along the horizontal plane with a vibratome to acquire 400 µm slices containing the hippocampus. The hippocampus was subdissected free from the rest of the slice and the CA3 subfield was removed. These slices were then quickly placed in a humidified holding chamber at room temperature (20-22°C) for 90 min recovery in recording ASCF (120mM NaCl, 3mM KCl, 1.5mM MgCl_2_, 1mM NaH_2_PO_4_, 2.5mM CaCl_2_, 26mM NaHCO_3_, and 10mM glucose) before starting recording experiments.

Schaffer collateral fibers were stimulated by placing a bipolar electrode (100 µs duration at 0.05 Hz; FHC, Bowdoin, ME) in the stratum radiatum of the CA1 subfield and an ACSF-filled glass recording pipette (3-5 MΩ; World Precision Instruments, Sarasota, FL) recorded a field excitatory postsynaptic potential (fEPSP) in the same layer of CA1. An input/output (I/O) curve was used to determine basal synaptic efficacy over a range of 10 stimulus intensities (25-300 µA) at an interstimulus interval of 20 s. The stimulus intensity was then modified to elicit 35-50% of the maximal fEPSP slope and paired-pulse fEPSPs (50 ms interpulse interval; PPR) were recorded each min for five min. Following paired-pulse recordings, individual stimulus pulses were applied every 20 s for 10 min, and baseline fEPSP responses were monitored. For bath application experiments, PPR was determined immediately after 35 min wash-out of VEH or HNK. For long-term potentiation (LTP) experiments, after recording baseline responses for 10 min, a high-frequency stimulation (HFS) protocol (4×100 Hz/1 s train at 20 s intervals) induced LTP, and fEPSP responses were monitored for the subsequent 60 min.

### Experimental Design and Statistical Analyses

Electrophysiology data were digitized at 10 kHz, filtered at 3 kHz, and analyzed with pCLAMP 10.7 software (Axon Instruments, Sunnyvale, CA). fEPSP values were normalized to the average fEPSP slope response recorded during the last five min of baseline. Individual normalized slope responses represent the average normalized slope value recorded at 20 s intervals successively over a one min period. Slices with an average baseline fEPSP slope value from 1-5 min that exhibited >10% variation compared to 6-10 min were excluded from the analysis. The effect of bath-applied vehicle/HNK or LTP magnitude was calculated by averaging the normalized fEPSP slope values collected during the last five min of recording. Reported n-values for in vitro drug application experiments indicate the number of slices assessed whereas ex vivo drug treatment experiments indicate the number of mice assessed. Within slice two-group comparisons of experiments with one factor with two levels were completed with paired two-tailed Student’s t tests whereas between-group comparisons of such experiments were conducted via two-tailed Student’s t-tests. Experiments that involved one factor with greater than two levels were analyzed via one-way analysis of variance (ANOVA). Between-group comparisons of experiments with more than one factor were analyzed via two-way ANOVA (e.g., factor A: HNK treatment; factor B: drug pretreatment or posttreatment). I/O curves were analyzed with one-way (factor A: HNK treatment) or two-way (factor A: HNK treatment; factor B: drug pretreatment or posttreatment) repeated measures ANOVA followed by Geisser-Greenhouse correction. Treatment means were separated via Holm-Šídák in instances of post-hoc comparisons when a significant interaction effect was detected. Data were assessed for normality (D’Agostino & Pearson) and homogeneity (Brown-Forsythe). In cases in which data were not normally distributed (i.e., Figs. 5D, 5K-L), data were log-transformed and analyzed via two-way ANOVA. Multiple linear regression analysis (least squares) was used to determine if the slope values depicting the relationship between PPR (independent variable) and the I/O response elicited at the maximal stimulus intensity (dependent variable) between slices treated with VEH or HNK were significantly different. Sample sizes were based upon our prior experience using similar paradigms. All data are presented as mean ± SEM. An *α* level of 0.05 was used as the criterion for statistical significance.

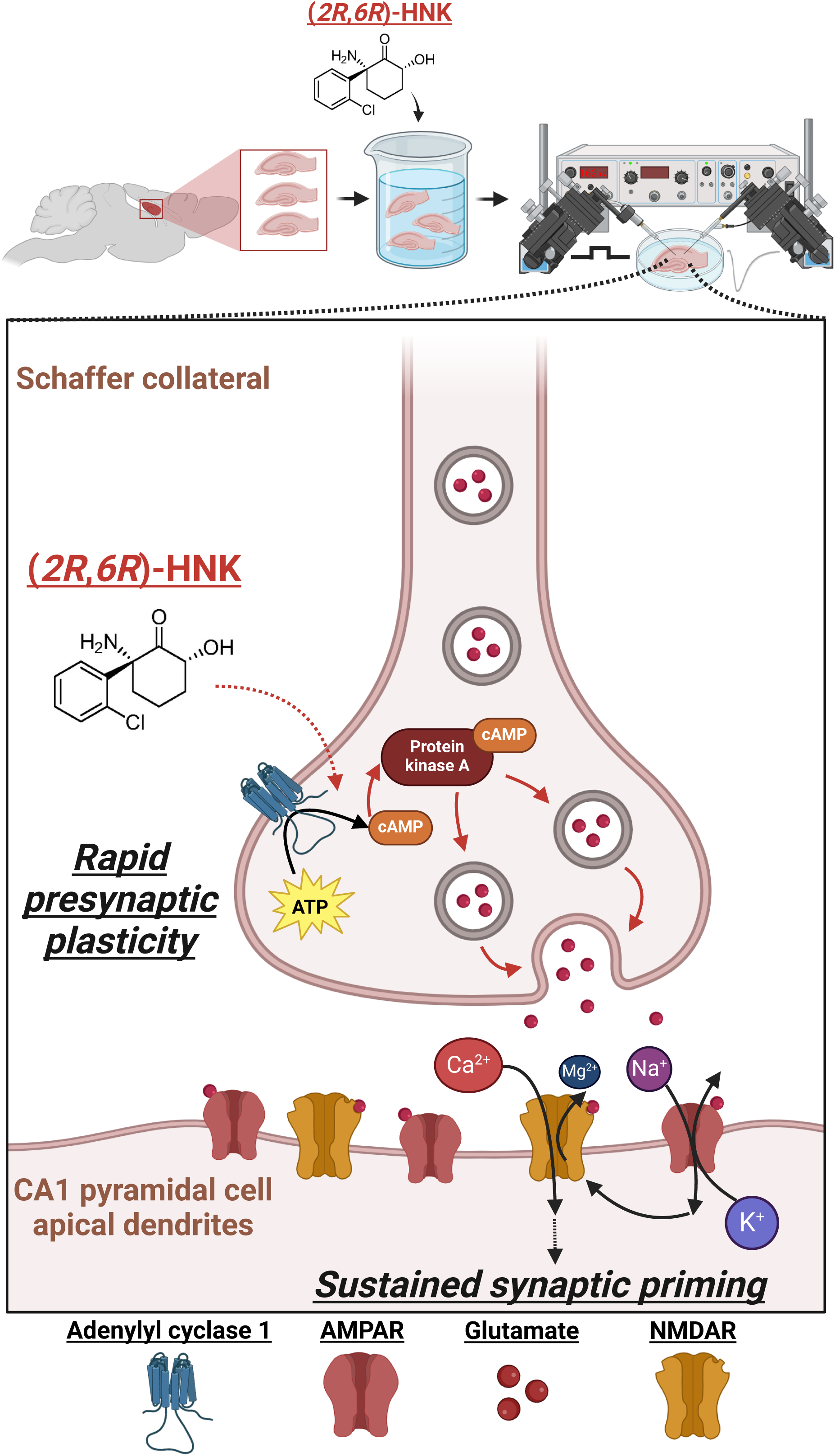

